# Detection and Molecular Characterization of Animal Adenovirus and Astrovirus from Western Maharashtra, India

**DOI:** 10.1101/2023.05.01.539015

**Authors:** PM Sawant, RB Waghchaure, PA Shinde, AP Palikondawar, M Lavania

## Abstract

**Background:** Astroviruses (AstV) and adenoviruses (AdV) are associated with diarrhoea in young ones of animals. However, the epidemiology and genetic diversity of AstVs and AdVs in animals are not well studied. Hence the present study was conducted to detect and characterize AstVs and AdVs in calves, piglets and puppies in India.

**Findings:** Out of the processed porcine (48), canine (80), and bovine (65) faecal samples from Western Maharashtra, India, the porcine AstV (PAstV), bovine AstV (BAstV), canine AstV (CAstV), and porcine AdV (PAdV) were detected in 12.5%, 7.69%, 3.75% and 4.1% of samples, respectively. In the RNA-dependent RNA polymerase region-based phylogenetic analysis the detected BAstV strains grouped with MAstV-28, MAstV-33, and MAstV-35, CAstV strains belonged to MAstV-5, PAstV strains belonged to MAstV-24, MAstV-26, and MAstV-31. Both the detected PAdV were of genotype 3, exhibiting 91.9-92.5% nucleotide identity with Ivoirian and Chinese strains. The study reports first-time BAstVs from calves and PAdV-3 from piglets in India.

**Conclusion:** The study revealed diversity in the circulation of AstVs and AdVs in young ones of dogs, pigs and calves. Therefore, extensive epidemiological investigations of AstV and AdV are necessary to confirm their association with diarrhoea and to develop diagnostic tools and control measures.

## 1. Introduction

Diarrhoea is the leading infectious cause of death among neonates of animals. The farmers and pet owners suffer a huge economic loss□from diarrheal disease. Therefore, monitoring of diarrhoeal diseases should be performed on a priority basis. Acute diarrhea is caused predominantly by rotavirus and norovirus worldwide. Some countries have introduced rotavirus vaccines in animals to provide effective Protection. However, studies on other diarrheal viruses such as AdV and AstV have been largely neglected.

AdVs are double-stranded DNA viruses belonging to the family *Adenoviridae*. Of the five serotypes of porcine adenoviruses (PAdV) belonging to Genus Mastadenovirus, PAdV-1-3, PAdV-4 and PAdV-5 correspond to three species PAdV-A, PAdV-B, and PAdV-C, respectively. Additionally, two PAdV serotypes□have also been proposed (Benkő et al., 2022). Although PAdVs rarely□cause diseases of economic concern in the domestic population. In deceased pigs, PAdV’s are found associated with certain diseases such as encephalitis, nephritis, respiratory diseases and reproductive disorders (Kumthip et al., 2019). The bovine adenoviruses (BAdV) belong to the genera, *Mastadenovirus* and *Atadenovirus*, with 10 recognized serotypes. The *Mastadenovirus* genus contains BAdV-1 (BAdV-A), BAdV-2 (Ovine adenovirus-B), BAdV-3 (BAdV-B), BAdV-9 (HAdV-C), BAdV-10 (BAdV-C) serotypes respectively whereas the serotypes BAdV-4, -5, -6, -7 and -8 are included in *Atadenovirus* genus as BAdV-D (Benkő et al., 2022). The majority of these serotypes cause mild diseases of the gastrointestinal or respiratory tract in the bovines (Gaba et al., 2019). Canine adenovirus (CAdV) belongs to the genus *Mastadenovirus*, divided into 2 serotypes CAdV-1 and CAdV-2 (Benkő et al., 2022). The CAdV-1 type causes infectious canine hepatitis, which is life-threatening in puppies and encephalitis in foxes, racoons, bears and skunks. The CAdV-2 type is a cause of kennel cough in breeding dogs (Gaba et al., 2019).

Astroviruses (AstVs) belonging to the family *Astroviridae* are non-enveloped, with single standard RNA as a genome arranged in three open reading frames; ORF1a and ORF1b codes for serine protease and RNA dependent RNA polymerase (RdRp), and ORF2 codes for capsid proteins. The family consist of 2 genera, *Mamastrovirus* which includes 19 species and *Avastrovirus* which includes 3 species.

The porcine AstVs (PAstVs) group into five lineages PAstV1-5 which have been classified into seven genotypes; MAstV-3, MAstV-22, MAstV-24, MAstV-26-27, MAstV-31-32. The bovine AstVs (BAstVs) are divided into seven genotypes MAstV-13, MAstV-28-30, and MAstV-33-35 while CAstVs are classified into MAstV-5 (Zhu et al., 2022; Bosch et al., 2011). The high genetic diversity and widespread presence of AstV make diagnosis and control strategies extremely difficult. Although AstV is found in the faeces of animals with and without diarrhea most studies reported the detection rate with symptoms of gastroenteritis is higher than a symptomatic one, indicating its potential role in diarrhoea.

Presently enzyme-linked immunosorbent assay and polymerase chain reaction (PCR) have enabled the detection of rotavirus in both animals and humans. However, surveillance and genetic characterization studies for animal AstV and AdV in India are limited. Therefore, the present study detected and partially characterized adenovirus and astrovirus from animal stool samples in the Western Maharashtra region of India.

## 2. Materials and methods

### 2.1 Sample collection

The stool samples from diarrheic and asymptomatic calves, piglets and puppies under six months of age were collected for the period 2017 to 2019 from Western Maharashtra, India. The faecal samples from 48 piglets (diarrheic 28; asymptomatic 20), 80 pups (diarrheic 55, asymptomatic 25), and 65 calves (55 diarrheic, 10 asymptomatic) were collected, transported on ice and stored at□-80°C until further processing. A 30% stool suspension was centrifuged at 10,000 rpm for 10 mins, supernatant was collected and processed for nucleic acid extraction.

### 2.2 Nucleic acid extraction

Total RNA and DNA were extracted from 140 μl of stool suspension using an RNA extraction kit (Qiagen, Germany), following the manufacturer’s recommendation. The extracted nucleic acids were stored at -80°C till their use as a template in reverse transcriptase polymerase reaction (RT-PCR) and PCR.

### 2.3 Conventional PCR for adenovirus detection

The conventional PCR for the detection of MastAdV and AtAdV used previously published primers (Sibley et al., 2011). The PCR reaction consisted of 5 μl of viral DNA, 1 μl of both forward and reverse primers, 0.5μl dNTPs, 1.5μl of 10X buffer, 0.75μl of 50mM MgCl_2_, 0.5μl *Taq* Polymerase enzyme (Invitrogen) and volume made to 25 μl with nuclease-free water (NFW). The thermocycling was carried out with an initial denaturation of 94°C for 4 min, thereafter 45 cycles of 94□ at 1 min, 50□ at 1 min, 72□ at 2 min and final extension for 7 min respectively.

### 2.3 □One-Step□□RT-PCR for astrovirus detection

The one-step RT-PCR used primers from a previously published panAstV assay which amplifies a 420bp fragment of the RdRp region (Chu et al., 2008). A 5μl of snap chilled RNA was added in 20μl of reaction mixture containing 3.5μl of NFW, 12.5μl of 1X reaction buffer, 1μl of SuperScript III RT/Platinum *Taq* Polymerase mix (Invitrogen), 1μl each of forward and reverse primer. The RNA was denatured at 95□ for 5min immediately followed by a snap chill at 4□ for 5min. After that cDNA was synthesized at 42□ for 45min, then denaturation at 95□ for 3 min, followed by 40 cycles of 94□ for 15sec, 50□ for 30 sec, 68□ for 2min, final extension step was performed at 68□ for 7 min respectively. The positive samples were visualized onto 1.5% agarose gel and observed by the Gel Documentation system (Bio-Rad, USA).

### 2.4 Sequencing and phylogenetic analysis

The purified PCR amplicons were subjected to cycle sequencing using Big Dye Terminator cycle sequencing kit v 3.1 (Applied Biosystems, CA, USA), and the dye terminators from the reaction were cleared with DyeEx 2.0 Spin Kit (Qiagen, Germany). After post-purification, the sequencing was performed on an automated Genetic Analyzer ABI-PRISM 3720 (Applied Biosystems, USA). The chromatograms were analyzed in Sequencing Analysis 5.2.0 (Applied Biosystems, USA). The BLAST was used for sequence similarity assessment. The study sequences and GenBank database sequences were aligned using CLUSTALW with the MEGAv6.06 programme. The Neighbour-joining phylogenetic trees were constructed using the Kimura-2 parameter method, and 1000 bootstraps were used to estimate the confidence of the branch. The nucleotide identity was estimated using the same software by pairwise distance using the p-distance model.

## 3. Results

### 3.1 Adenovirus detection and phylogenetic analysis

A total of two (4.1%) porcine stool samples were positive for PAdV, while canine and bovine samples were negative for CAdV, and BAdV. The partial hexon gene of detected strains shared 93.4% of identity between themselves. The PAdV study strain exhibited 91.9-92.5% identity with PAdV-3 strain PGOU244/Cote d’Ivoire/2012 and Chinese PAdv-SCMS01 (Fig 1). However, the study strains exhibited 64.8–66% identity with PAdV-5 strains. The curated sequences were deposited in GenBank (OQ852903 and OQ852904).

**Fig.1.**
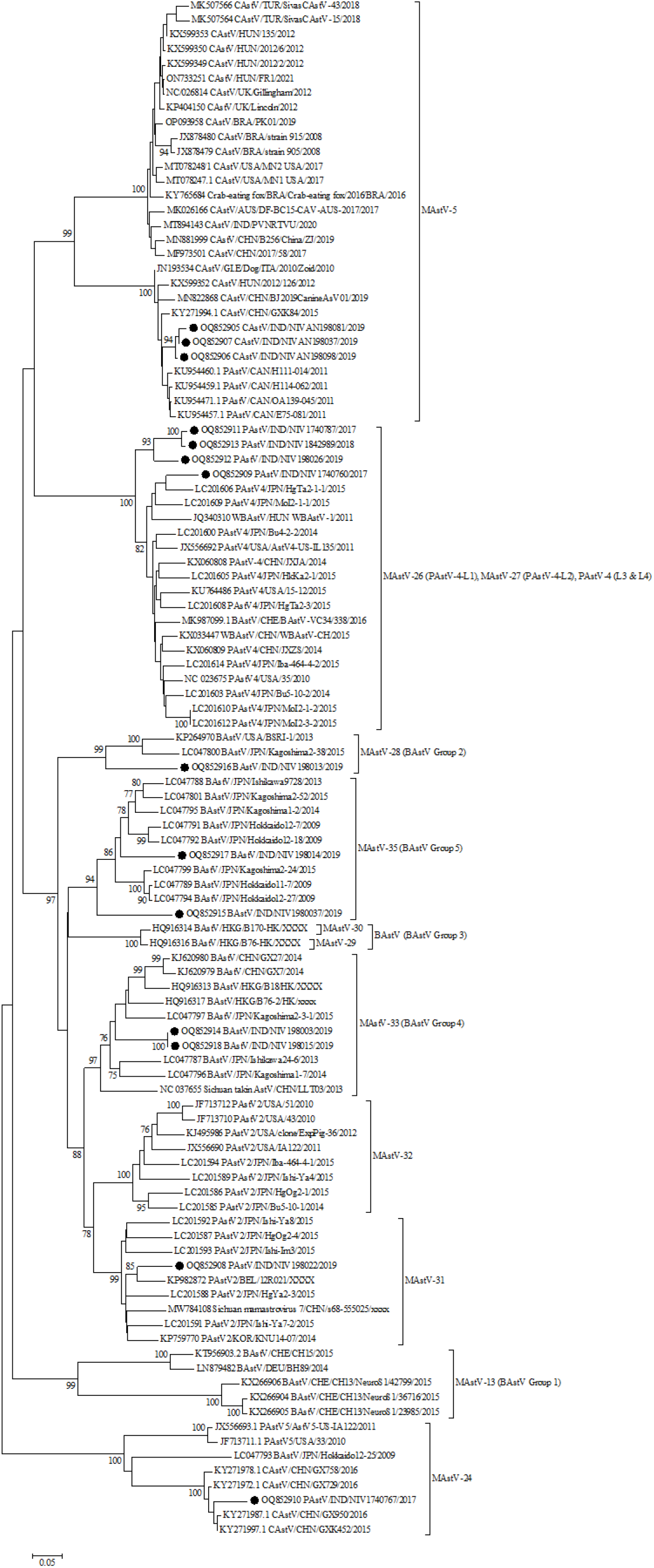
Phylogenetic analysis of RdRp nucleotide sequences of astroviruses detected from porcine, canine, and bovine stool and faecal samples. The sequences detected in the present study are marked with a black solid circle.

### 3.2 Astrovirus detection and phylogenetic analysis

The RdRp region nucleotide sequences of six PAstV, five BAstV and three CAstV study strains were compared with reference sequences from GenBank. The PAstV strains (NIV1740760, NIV1740787, NIV198026, and NIV1842989) displayed 88.1% identity with Chinese PAstV4 wild boar strain WBAstV-CH (MAstV-26) (Fig 2). The remaining two study strains, NIV198022 showed 90.3% identity with PAstV2 strain Bel-12R021 (MAstV-31), and NIV1740767 showed 76.3-76.6% identity with American PAstV5 (MAstV-24) strains AstV5-US-IA122 and USA/33. The detected CAstV strains, NIVAN198081, NIVAN198098, and NIVAN198037 were identical and exhibited maximum identity in the 93.7- 95.4% range with Chinese BJ2019 CanineAsV01 and HUN/2012/126 (MAstV-5). All the detected BAstV strains shared nucleotide identity between 66.3-71%. The NIV198003 and NIV198015 (G4) were identical and displayed 86.2% identity with JPN/Kagoshima2-3-1 (MAstV-33), and study strain NIV198014 exhibited 87.7-87.4% identity with BoAstV/JPN/Hokkaido12-18 and BoAstV/JPN/Hokkaido12-27 (MAstV-35), study strain NIV1980037 (G5) with (80.5%) BoAstV/JPN/Kagoshima2-24 and BoAstV/JPN/Hokkaido12-27 (MAstV-35), NIV198013 (G2) (79.7%) with USA/BSRI-1 and BoAstV/JPN/Kagoshima2-38 (MAstV-28). All the curated sequences were deposited in Gen Bank with accession numbers (OQ852905-OQ852918).

**Fig.2.**
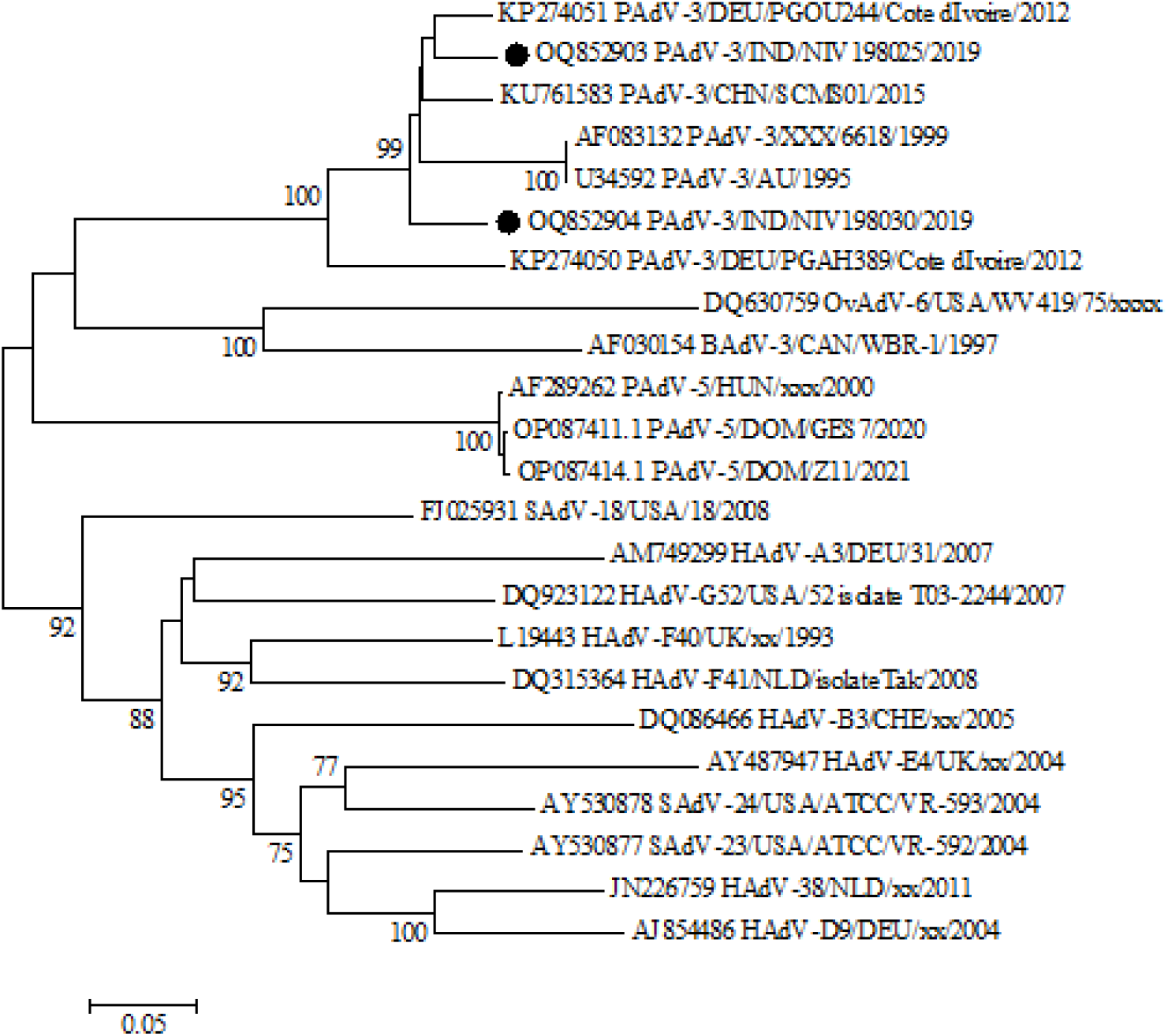
Phylogenetic analysis of RdRp nucleotide sequences of adenoviruses detected from the porcine stool and faecal samples. The sequences detected in the present study are marked with a black solid circle.

## 4. Discussion

The enteric viruses, astroviruses and adenoviruses, are considered to be associated with diarrhoea in various animal species including pigs, dogs, cats, calves, sheep and goats etc. The present study on screening of dog, pig and bovine samples revealed 12.5% of PAstV, 7.69% of BAstV, 3.75% of CAstV and 4.1% of PAdV.

The PAdV are suspected to be widespread in pigs, and among five PAdVs, PAdV3 is the most prevalent and well-characterized serotype (Mittal et al., 2016); however, their detection has not been reported from pigs reared in India. The present study identified and retrieved a total of two adenovirus sequences from piglets. After analysis of the partial hexon gene, the study strains found similarity with PAdV-3 reported from Africa (Cote d’Ivoire) and Asia (China) with 91.9-92.5% nucleotide identity. Our study reports the first detection of PAdV-3 from the Indian pig population, indicating that the PAdV3 circulating in Asia and Africa are genetically similar. The properties of PAdV-3 like genetic and structural similarities with HAdV-5, no cross-neutralization with Anti-HAdV-5 antibodies and transduction of human cells, imply that the development of PAdV-3 vectors will be a supplement to HAdV-5 vectors (Mittal et al., 2016).

Five genotypes of PAstV (PAstV1-5) have been identified with varying prevalences from both healthy and diarrheic pigs, including in the Czech Republic, Canada, Columbia, USA, Hungary, China, South Korea and Thailand (De Benedictis et al., 2011; Kumthip et al., 2018; Rawal and Linhares 2022).

However, knowledge of PAstV diversity in Indian pigs is limited and only two studies have reported genotypes (Kattoor et al., 2019; Kour et al., 2021). One study revealed the circulation of PAstV1, PAstV2, PAstV4, and PAstV5 in Haryana state with a predominance of the PAstV1 genotype. The second study covering a wider geographic area reported a predominance of PAstV4 and the detection of PAstV2 genotypes. In the present study, one porcine study strain clustered with American PAstV5 strains, including BoAstV/JPN/Hokkaido12-25/2009 (LC047793.1), believed to be derived from cross-species transmission (Nagai et al., 2015). The prevalence of PAstV (10.41%) in the present study was lower as compared to all published reports irrespective of the RT-PCR method used except one Chinese study which reported a 2.82% positivity employing sequence-independent-single primer amplification (SISPA) (Li et al., 2015; Rawal and Linhares, 2022).

Canine AstV has been reported to be associated with acute gastroenteritis or without clinical symptoms across the world. The only recently published study from the Hyderabad region of India reported a 7.2% prevalence of CAstV, and the strain with full genome sequence grouped with Chinese and Brazilian strains (Dema et al., 2023). In contrast, all the present study strains are grouped with a different cluster of Hungarian, Italian and Chinese strains. The studied strains were from stray dogs which were maintained along with abandoned foreign dog breeds maintained in shelters in Pune city. Hence the circulation of CAstVs resembling Hungarian, Italian and Chinese strains is quite possible because of the ingestion of faecal material from foreign breeds. This suggests that there is a higher diversity of CAstV in India.

Most of the BAstVs detected in faecal samples from North and South America, Asia, Europe & African regions (Egypt) fall into G1, G2, G3, G4 and G5 groups (Zhu et al., 2022). The sequencing of the partial RdRp gene in the present study revealed the presence of BAstV from divergent lineages G2 (MAstV-28), G4 (MAstV-33), and G5 (MAstV-35). In the phylogenetic tree all but one strain clustered with Japanese bovine strains. The NIV198013 grouped with American and Japanese bovine strains. Importantly, no study strain found clustering with strains reported from neurological disease. The present study reported a much lower prevalence which is in concurrence with the Korean and the United

Kingdom study (Oem and An, 2014; Sharp et al., 2015). Nevertheless, to our knowledge, the present study reports the first-time detection of bovine AstVs from India. The majority of AstV sequences are fragments of length 300-400bp which is not appropriate to study cross-species transmission and recombination, hence complete genome sequencing is suggested to study the evolution of AstVs.

In conclusion, the study found AstVs circulating in puppies, piglets and calves with significant genetic diversity. However, AdV was detected only in piglets with identity matching with strains from diverse geographical regions. The scarce genetic data of AstVs and AdV from the Indian animal population will challenge the laboratory diagnosis and prophylaxis strategies. In the future, large-scale surveillance targeting astrovirus and adenovirus should be initiated to understand genetic diversity, its contribution to disease, and the eventual exploitation of adenovirus as a xenogeneic vector for gene therapy and vaccine delivery.

## Author contributions

Conceptualization, PMS,; Sample Collection, PMS; Sample Screening, RBW, PAS, PMS,; Data analysis, PMS, APP, ML,; Manuscript Writing & editing, PMS, ML, APP.

